# Multi-scale Loading and Damage Mechanisms of Rat Tail, Plantaris, and Achilles Tendons

**DOI:** 10.1101/471052

**Authors:** Andrea H. Lee, Dawn M. Elliott

## Abstract

Tendinopathy, degeneration of tendon that leads to pain and dysfunction, is common in both sports and occupational settings, but multi-scale mechanisms for tendinopathy are still unknown. We recently showed that micro-scale sliding (shear) is responsible for both load transfer and damage mechanisms in rat tail tendon; however, rat tail tendon is a specialized non-load bearing tendon, and thus the load transfer and damage mechanisms are still unknown for load-bearing tendons. The objective of this study was to investigate the load transfer and damage mechanisms of load-bearing tendons using rat plantaris and Achilles tendons. We demonstrated that the micro-scale sliding is a key component for both mechanisms in plantaris tendon, similar to tail tendon. Namely, the micro-scale sliding was correlated with applied strain, demonstrating that load was transferred via micro-scale sliding in the plantaris and tail tendons. In addition, while the micro-scale strain fully recovered, the micro-scale sliding was non-recoverable and strain-dependent, and correlated with a tissue-scale mechanical parameters. When the applied strain was normalized, the % magnitudes of non-recoverable sliding was similar between the plantaris and tail tendons. Achilles tendon demonstrated some of the mechanical responses observed in plantaris and tail tendons, yet the results were inconclusive due to its complex structure. Statement of Clinical Significance: Understanding the mechanisms responsible for the pathogenesis and progression of tendinopathy can improve prevention and rehabilitation strategies and guide therapies and design of engineered constructs.

## Introduction

Tendinopathy, degeneration of tendon that leads to pain and dysfunction, is common in both sports and occupational settings,^1^ but the mechanisms for tendinopathy are still unknown. The mechanisms of tendinopathy have been attributed to overloading that leads to inferior mechanical and microstructural degradation,^2–5^ and these occur across multiple hierarchical scales.^6–12^ Rodent tendon models of overloading have frequently been used to study tendinopathy; ^4,5,13–16^ however, the multi-scale mechanisms of normal load transfer and damage are not well understood.

In order to measure multi-scale load transfer and damage mechanisms, where damage is defined as an irreversible change at the micro-scale that is observable in the tissue-scale mechanics, the structure and mechanics must be simultaneously measured across multiple length scales. At the micro-scale, there is evidence that under high or repeated tensile loading, microstructural changes occur that may represent damage.^15,17,18^ At the tissue-scale, separate studies have provided evidence that mechanical properties, such as reduced modulus or peak stress, change under loading, which may represent damage.^5,18,19^ In addition, a significant advancement in tendon damage modeling highlights the need to study damage across multiscales.^20–25^ Still, multi-scale load transfer and damage mechanisms of tendons are not fully elucidated because simultaneous quantification of the microstructural changes and tissue-scale mechanical properties is lacking.

Using rat tail tendon, we recently showed that micro-scale sliding is a mechanism for load transfer,^14^ and non-recoverable micro-scale sliding in response to high tensile loading leads to tendon damage.^4^ Yet, rat tail tendon is specialized and is a relatively non-load bearing tendon,^10,26^ and thus the load transfer and damage mechanisms are still unknown for load-bearing rat tendons. Despite some differences between rat tail tendon and load-bearing tendons, we hypothesize that micro-scale sliding is a mechanism for both tendons since they both have similar fiber structures.^10^ We chose rat plantaris and Achilles tendons to study these mechanisms. The plantaris tendon bears high-stress *in vivo*^27^ and has a relatively simple structure.^10^ The Achilles tendon also bears high-stress and is a highly specialized tendon with a complex structure, where three sub-tendons fuse together to form a single tightly bound tendon.^10,28–31^

The objective of this study was to investigate the multi-scale load transfer and damage mechanisms of load-bearing tendons, rat plantaris and Achilles tendons, and to evaluate if the mechanisms are similar to the mechanisms demonstrated in rat tail tendon. Recently published data from the tail tendon, which has the simplest structure,^10^ was reanalyzed and included for comparison.^4^ We hypothesize that the micro-scale sliding will increase with applied strain and will be non-recoverable when loaded to high strain in the plantaris and Achilles tendons.

## Methods

Mechanical testing was performed on rat plantaris (n=33) and Achilles (n=6) tendons that were harvested from 4-7-month-old female Long Evans rats. Additionally, the mechanical testing data of rat tail tendon (n=28) from 6-8-month-old male Sprague-Dawley rats were reanalyzed for this study.^4^ All tendons had been frozen at −20C, and the maximum number of the freeze-thaw cycle was limited to three to minimize the effect of freezing.^32^ A total summary of sample number and dimensions, including cross-sectional-area and gauge length, are included as a supplemental material (Fig S-1).

### 1. Sample Preparation

A plantaris and Achilles tendon complex was dissected from a calcaneus bone, and each tendon was separated as shown previously.^10^ The rat tail tendon was harvested as previously described.^32^ All tendons were placed in PBS at room temperature to keep hydrated. We stained rat plantaris and tail tendons with 10 *μ*g/ml 5-DTAF (5-(4,6-Dichlorotriazinyl) aminofluorescein, Life Technologies) and Achilles tendon with 20 *μ*g/ml 5-DTAF for 20 minutes.^33^ We mechanically tested tendons on a custom-made uniaxial testing device with a PBS bath mounted on an inverted confocal microscope (LSM 5 LIVE, objective Plan-Apochromat 10x/0.45) as previously described.^14^

### 2. Mechanical Analysis

#### 2.1 Mechanical Testing Protocol

Plantaris tendons were randomly assigned to 3 strain groups with n=11 tendons per group, where each group was mechanically loaded to a grip strain of 14, 20, or 25%. Each Achilles tendon was loaded to a single grip strain of 14%, with n=6 tendons. After mechanically testing one strain group of Achilles tendon, we decided to discontinue testing other strain groups due to the complexity of the structure^10^ and a related limitation in our testing setup (Fig 1). The details will be further elaborated in future sections.

**Fig 1:**
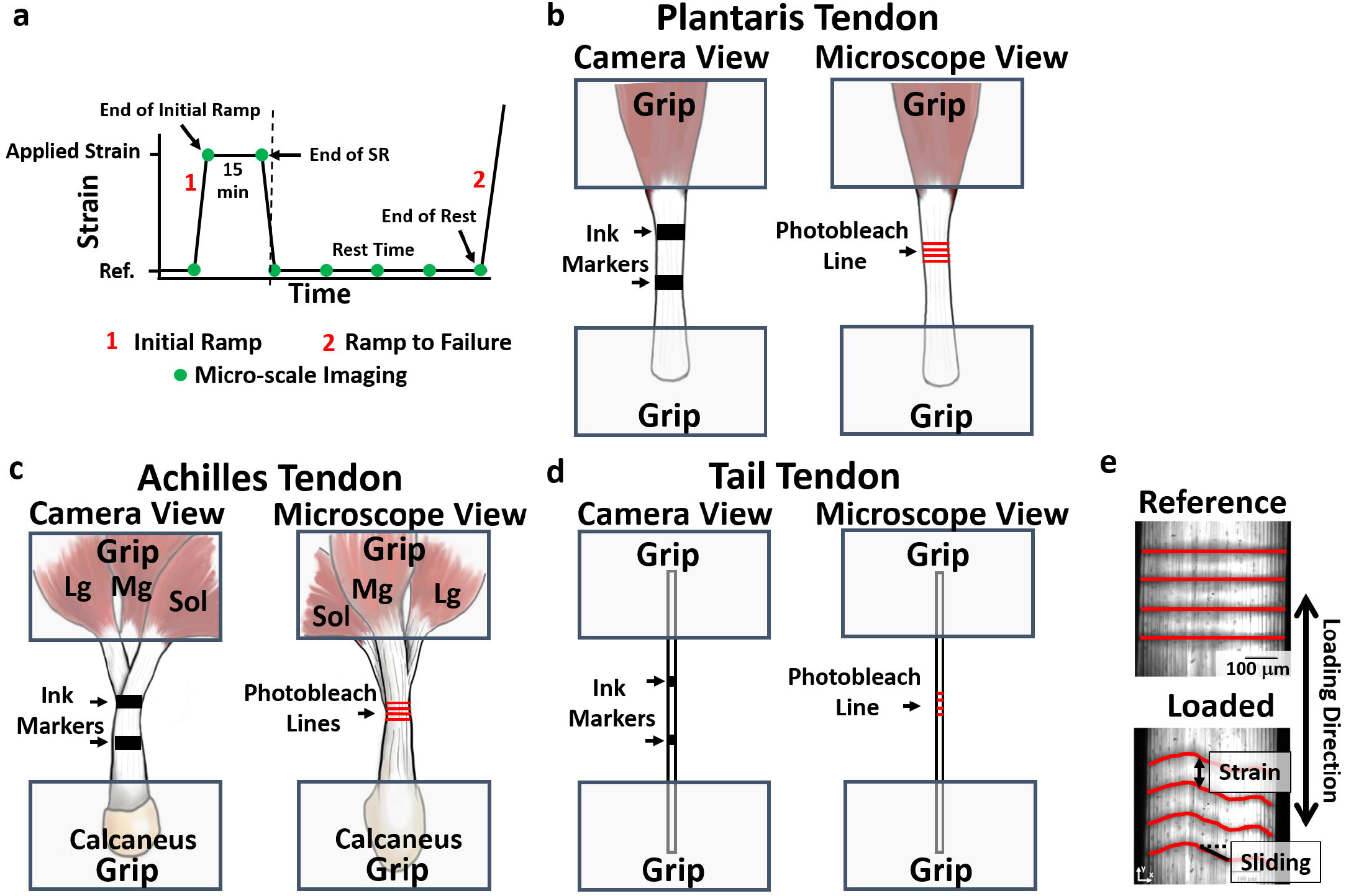
Schematic of experimental design and overall micro-scale deformation. a) A schematic of mechanical testing protocol is shown. We quantified tissue-scale parameters at two time points, the Initial Ramp (1) and Ramp to Failure (2), and micro-scale parameters at ten time points shown as micro-scale imaging (green dot). The parameters measured at End of Initial Ramp, End of Stress Relaxation (SR), and End of Rest were quantified and used for further analysis. Dashed line represents when tendon was unloaded to the reference length. b) Ink markers were applied to plantaris tendon to optically measure the applied strain using a CCD camera. On the other side of the tendon, four photobleached lines were applied using a confocal microscope to measure the micro-scale deformations. Similarly, the ink markers and photobleached lines were applied for c) Achilles and d) tail tendons. The photobleach lines are not to scale and enlarged for visualization. e) The micro-scale strain and sliding, a measurement of micro-scale deformation of averaged fibril and fiber structures, were quantified using deformed photobleached lines.

We preloaded each tendon to 5mN to establish a reference length and preconditioned to 5 cycles of 3% grip strain. Each tendon was mechanically loaded to the grip strain, and we held the strain constant for 15 minutes of stress relaxation followed by unloading to the reference length (Fig 1a). The dashed line represents when tendon was unloaded. The tendon was held at the reference length for 40 minutes of rest period before ramping to failure. All loading and unloading strain rates were 1%/s.

The tail tendon was tested following a similar mechanical testing protocol as above.^4^ The tail tendon was randomly assigned to 4 strain groups with n=7 tendons per group, where each group was mechanically loaded to a grip strain of 2, 4, 6, or 8%. The tail tendon was preconditioned to 5 cycles of 4% grip strain and held at the reference length for 60 minute before ramping to failure.^4^

#### 2.2 Data Acquisition

To calculate the cross-sectional area, we used the confocal microscope and a scanning laser displacement sensor.^34^ We collected confocal image stacks under preload and fitted with an ellipse to measure a cross-sectional area. For the plantaris and Achilles tendons, only a major axis was measured with the confocal microscope, and a scanning laser displacement sensor was used to measure a minor axis to calculate the area. For the tail tendon, both major and minor axes were measured with the confocal microscope.

To measure the tissue strain, we applied two ink markers directly on the tendon using a permanent marker (Fig 1b-d). The ink markers were imaged with CCD camera to track displacement of markers using digital image correlation (Vic 2D, Correlated Solutions). The maximum tissue strain, measured at the End of Initial Ramp, was optically calculated and is labeled as ‘applied strain’. We applied ink markers on the posterior side of the plantaris tendon and on the soleus (Sol) sub-tendon for the Achilles tendon (Fig 1c). For the tail tendon, we applied ink markers on the mid-substance of the tendon, away from the grips (Fig 1d).

We quantified the micro-scale parameters (i.e., micro-scale strain and sliding) using the confocal microscope as previously described (Fig 1e, S-2).^4,14^ Four lines were photobleached using the confocal microscope, separated by 200μm at the mid-substance of tendons, (Fig 1b-e). We bleached lines on the anterior side of plantaris tendon and medial gastrocnemius (Mg) sub-tendon for the Achilles tendon (Fig 1c). We acquired confocal image stacks at 10 times points, represented by green dots (Fig 1a) The parameters measured at three time points, End of the Initial Ramp, End of Stress Relaxation (SR), and End of Rest, were primarily used for data analysis.

### 3. Data Analysis

#### 3.1 Tissue-scale Mechanical Parameters

The tissue-scale mechanical parameters were quantified at 2 time points: Initial Ramp and Ramp to Failure (Fig 1a). Representative stress-strain (*σ-ε*) responses of the three tendons are shown (Fig 2). The tissue-scale parameters were analyzed using the methods and procedure from our previously published paper.^4^ Briefly, an inflection point, defined as a point where the *σ-ε* curve shifts from strain-stiffening to strain-softening, was determined (Fig 2).^4,34^ The *σ-ε* response up to inflection point was used to determine the transition point and linear region modulus by fitting a nonlinear constitutive model.^4,35^ All the parameters calculated for this study are marked on the representative *σ-ε* of plantaris tendon (Fig 2), which includes transition strain and stress, linear region modulus, and inflection point strain and stress. The parameters were calculated for both Initial Ramp and Ramp to Failure. This paper will focus on three tissue-scale parameters, Ramp to Failure transition strain, inflection point strain, and linear region modulus, because these parameters have been shown to represent damage well.^4^

**Fig 2:**
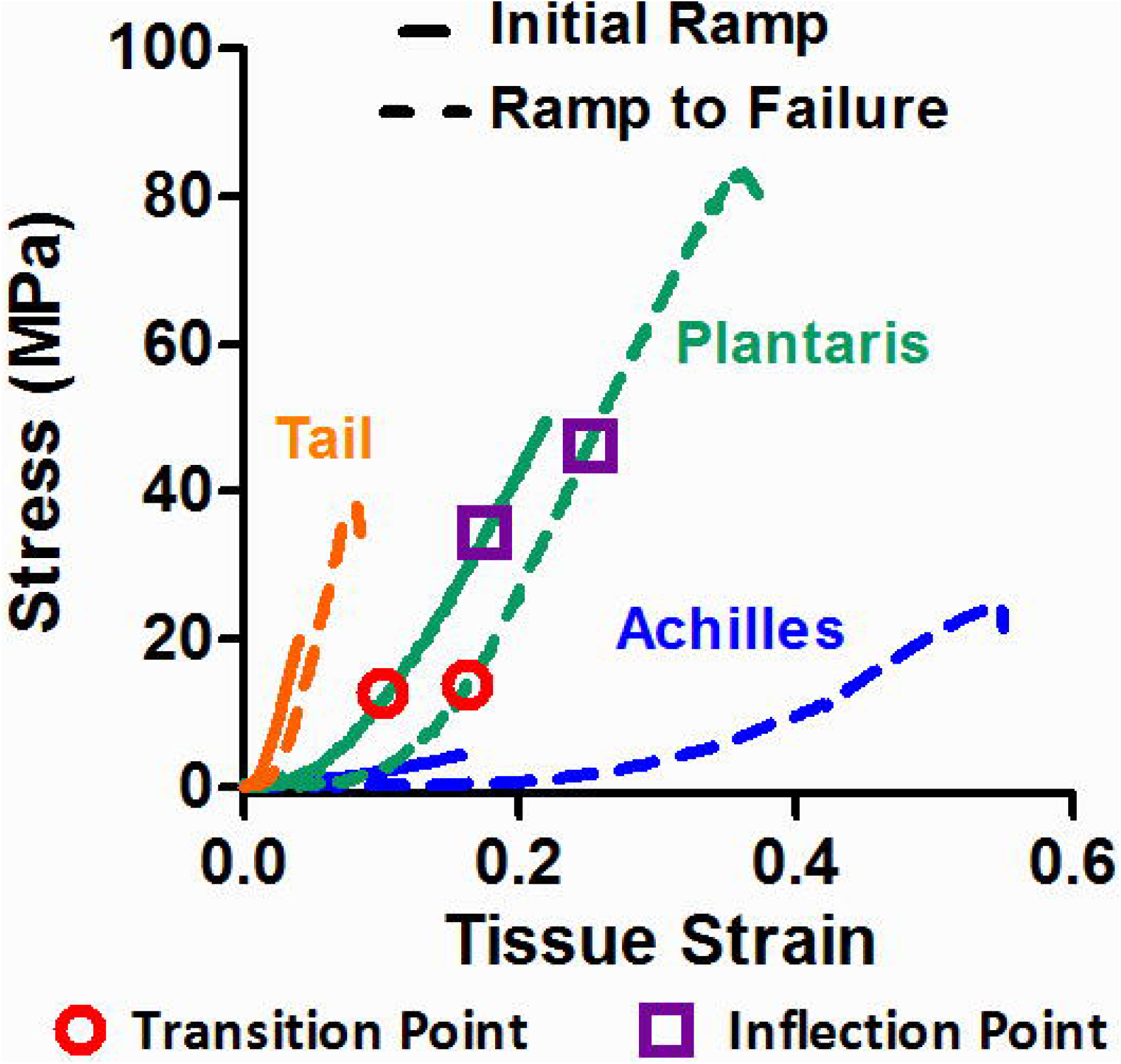
Tissue-scale parameter quantification. a) The representative *σ-ε* responses of plantaris, Achilles, and tail tendons are shown, where Initial Ramp is represented as a solid line and Ramp to Failure is represented as a dotted line. The stress relaxation and unloading are not shown in the graph. Transition point (red open circle), inflection point (purple open square), and linear region modulus (linear line between transition and inflection points) were quantified for both ramps. The representative sample for the plantaris tendon was loaded to 20%, Achilles tendon was loaded to 14%, and tail tendon was loaded to 6%.

The peak stress and strain were also quantified for all the tendons and can be found as a supplemental material (Fig S-3). In addition, the normalized applied strain was quantified by taking a ratio between averaged peak strain and applied strain of each sample (Fig S-3) to visualize all tendons on the same x-axis. The peak strain was not statistically different among strain groups for the plantaris and tail tendons (p > 0.2), and thus an averaged peak strain across all of the strain groups for each tendon was used. Many samples did not fail within the load cell limit and some tendons had complex failures, where failure initiated near or inside the grips. Hence, we decided to include peak strain and stress as a supplemental material, and the data should be used with caution.

#### 3.2 Micro-scale Parameters

To measure the micro-scale deformation, we used the deformation of photobleached lines.^14^ The micro-scale strain was defined as a change in distance between pairs of photobleached lines, and the micro-scale sliding was defined as averaged tortuosity (i.e., waviness) of all four lines (Fig 1e, S-2). The micro-scale strain represents an axial deformation, and the micro-scale sliding represents a relative shear deformation of collagen fibers and fibrils. The micro-scale strain and sliding were quantified for every time point (green dots). In addition, the ratio of micro: tissue strain ratio was calculated for the End of Initial Ramp to measure a strain attenuation from the tissue-scale to micro-scale (Fig 1a). This was calculated by taking a ratio between micro-scale strain and applied strain. The non-recoverable sliding, which represents damage at the micro-scale, was calculated using the following equation: 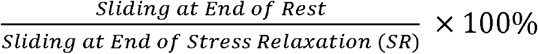. For visualization, only the first 40 minutes of the rest period was plotted for the rat tail micro-scale strain and sliding to match the rest period of plantaris and Achilles tendons, but the entire data from the rest period was used for the analyses^4^

#### 3.3 Excluded Samples

For the tissue-scale analyses, a total of two tendons were removed because one of the tail tendon fascicles was twisted and one of the plantaris tendons had a major grip slippage during the Initial Ramp. In these cases, the linear region modulus and applied strain were two standard deviations away from the population average. All Initial Ramp (Fig S-4) and Ramp to Failure parameters (Fig S-5) including transition and inflection point stress correlations are reported in supplemental material. For the micro-scale analyses, a total of eight samples across all groups were removed because photobleach lines were not clear enough to be recognized with an automatic registration. The final sample count can be found in the supplemental material (S.1).

### 4. Statistical Methods

Pearson’s correlations (r) with significance set at p=0.05 were conducted using GraphPad Prism. To determine which tissue-scale parameters show damage associated changes and depend on the applied strain, each of the Ramp to Failure tissue-scale parameters was linearly correlated with the applied strain. To study micro-scale deformation and determine the loading mechanism, micro-scale strain, micro: tissue strain ratio, and micro-scale sliding measured at End of Initial Ramp was linearly correlated with the applied strain. To measure the micro-scale damage and determine if it is dependent on the applied strain, we linearly correlated the micro-scale sliding with the applied strain. In addition, to determine the damage mechanisms, the micro-scale non-recoverable sliding was correlated with the applied and normalized target strains. To relate the damage at the micro-scale to the changes of mechanical response at the tissue-scale, the non-recoverable sliding was correlated with the Ramp to Failure properties.

## Results

### Tissue-scale Mechanical Parameters and Damage

The *σ-ε* curve of each tendon has unique characteristics, likely due to the structural differences among these tendons (Fig 2).^10^ The tail tendon bears the lowest strain, the plantaris tendon bears the highest stress, and the Achilles tendon bears the lowest stress but the highest strains among three tendons. Notably, the maximum applied strain was 8% for the tail tendon, whereas the maximum applied strain was 25% for the plantaris tendon. These strains are about 80% of the peak failure strain (Fig S-3), motivating the choices in strain groups based on our pilot test. The grip strain was within the standard deviation of the optically measured applied strain for all tendons, confirming the *σ-ε* curve was not a result of a premature failure or grip slippage (Fig S-4).

We quantified the Ramp to Failure mechanical properties (transition strain, inflection point strain, linear region modulus) from the *σ-ε* curves to investigate if damage was observed in the mechanical properties and, therefore, is strain-dependent (Fig 3). The transition strain (Fig 3a, p<0.0001), inflection point strain (Fig 3d, p<0.001), and linear region modulus (Fig 3g, p<0.05) significantly correlated with the applied strain for the plantaris tendons but not for the Achilles tendon (Fig 3b, e, h). This demonstrates that these parameters show damage that is strain-dependent in the plantaris tendon, similar to our previous finding in the tail tendon (Fig 3c, f, i).^4^

**Fig 3:**
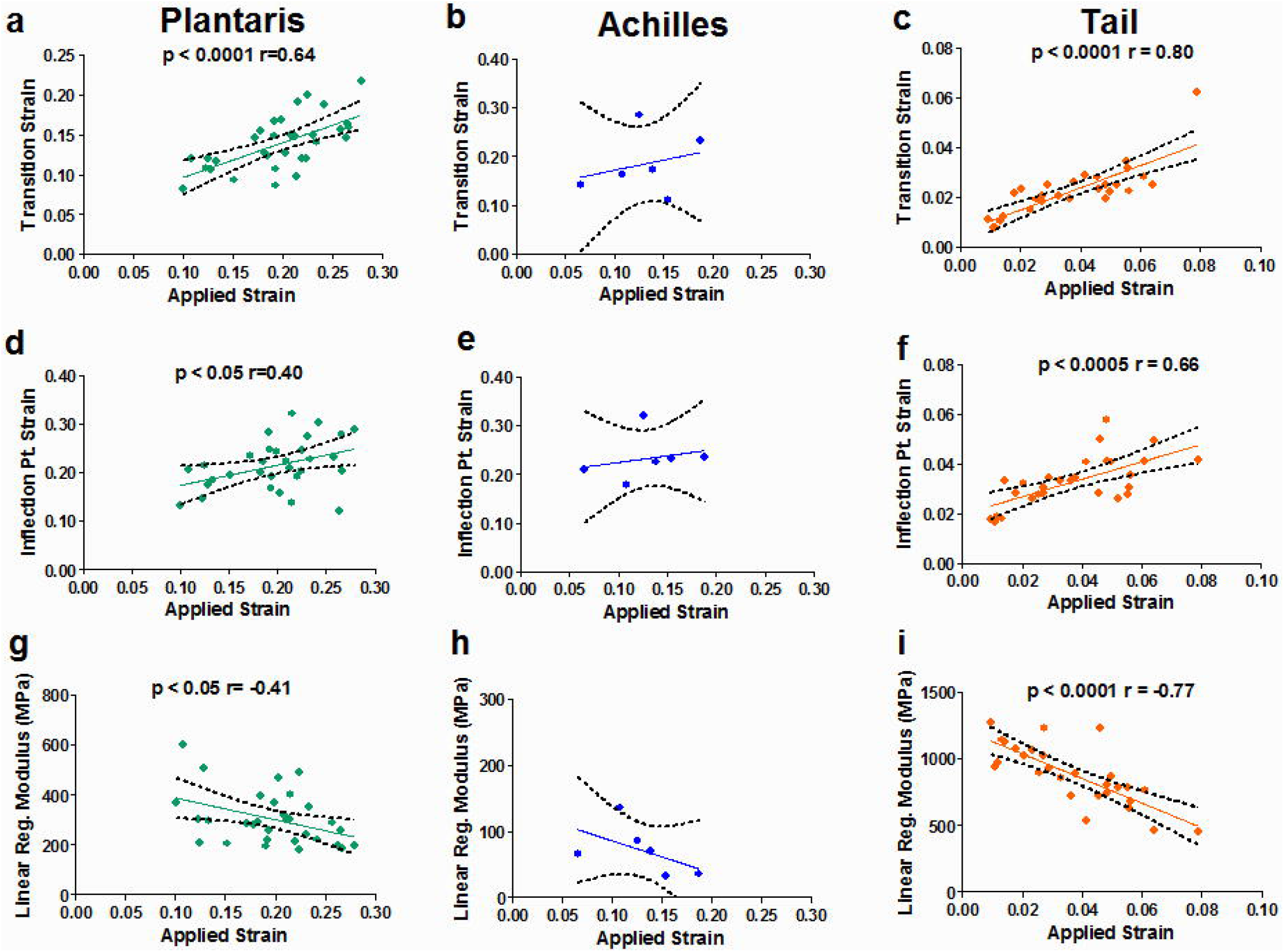
Tissue-scale mechanics of Ramp to Failure parameters. a)-c) The transition point strain, d)-f) inflection point strain, and g)-i) linear region modulus of plantaris, Achilles, and tail tendons were linearly correlated with the applied strain. The correlations of transition strain, inflection point strain, and linear region modulus of both plantaris and tail tendons were statistically significant, demonstrating that these parameters show damage and depend on the applied strain. None of the parameter of Achilles tendon correlated with the applied strain. Dashed lines represent 95% confidence intervals.

None of the parameters for Achilles tendon correlated with the applied strain, and this is likely due to the complexity of the structure.^10^ The tissue-scale parameters were quantified on the soleus sub-tendon, while the micro-scale parameters were quantified at medial gastrocnemius sub-tendon (Fig. 1d), and thus each tissue- and micro-scale parameter was measured at different sub-tendons. This mismatch of tissue- and micro-scale quantification is likely the cause of the observed inhomegenaity. Indeed, both human and rat Achilles tendons experience inhomogeneous strain.^30,31,36^ Thus, we discontinued testing of Achilles tendon after the 6 samples loaded to 14% applied strain.

### Multi-scale Tendon Loading Mechanisms

To study multi-scale deformation during loading, the micro-scale strain, micro: tissue strain ratio, and micro-scale sliding (quantified at the End of Initial Ramp) were correlated with the applied strain (Fig 4). The micro-scale strain did not correlate with the applied strain for the plantaris and Achilles tendons (Fig 4a-b), in contrast to the high correlation previously observed in the tail tendon (Fig 4c).^14^ The micro: tissue strain ratios (micro-scale strain divided by applied strain) were less than 1 and decreased with increasing applied strain for all tendons as expected (Fig 4d-f, p<0.05), demonstrating that strain is attenuated and the amount of strain attenuation is dependent on the applied strain. Notably, this ratio was much lower in the plantaris and Achilles (~0.2) compared to the tail (~0.5-1.0). The micro-scale sliding of the plantaris and tail tendons increased with increasing applied strain (Fig 4g-i, p <0.05). The magnitude of the sliding of the Achilles tendon was low, and the sliding did not correlate with the applied strain. Despite the differences in applied strains and magnitudes of the responses, the directions of the correlations were the same between the plantaris and tail tendons, whereas those of Achilles tendon were different.

**Fig 4:**
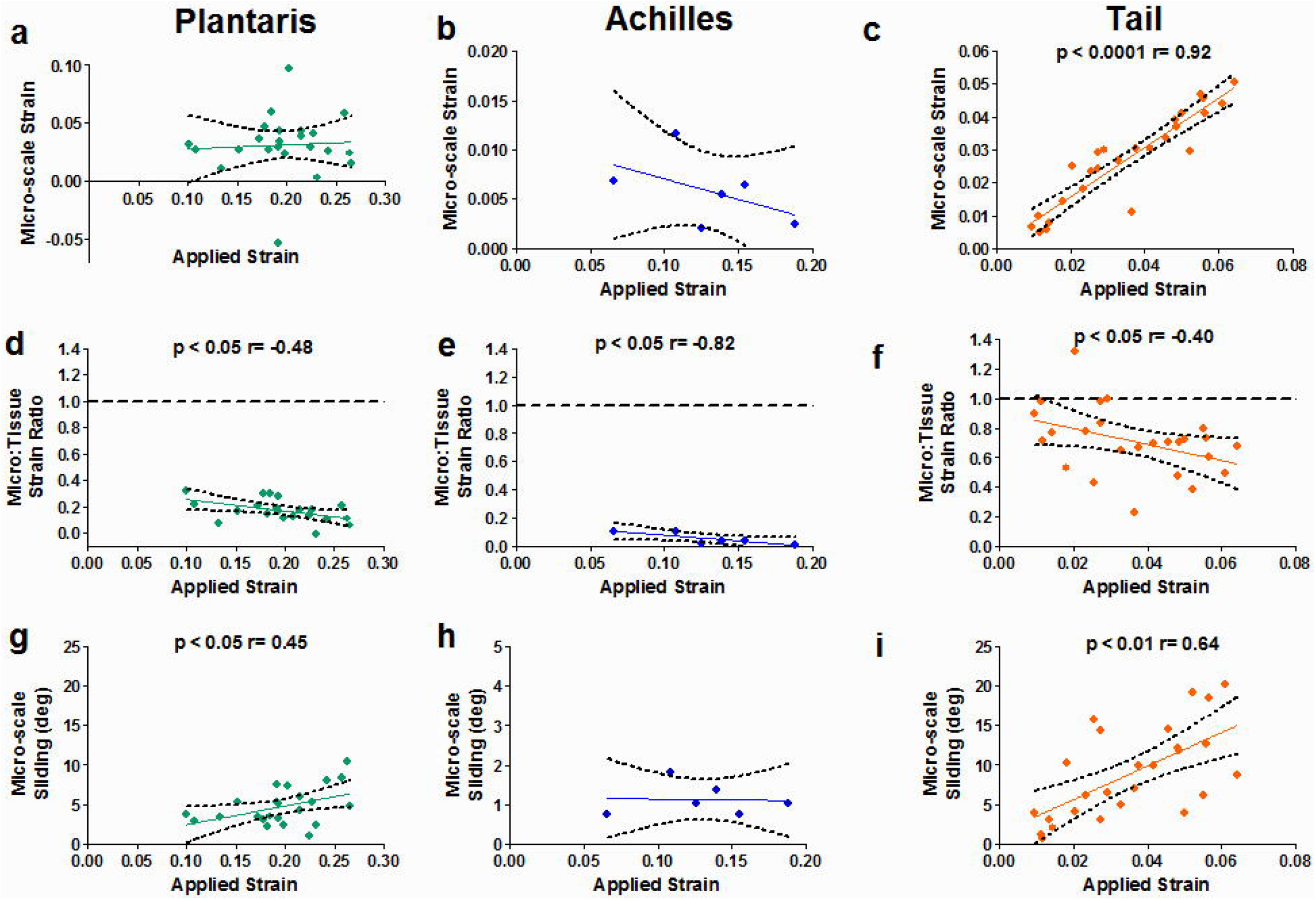
Loading mechanisms at multi-scale measured at the End of Initial Ramp. a)-c) The micro-scale strain, d)-f) micro: tissue strain ratio, and g)-i) micro-scale sliding of plantaris, Achilles, and tail tendons were linearly correlated with the applied strain. The micro-scale strain increased with the applied strain for the tail tendon, but not for the plantaris and Achilles tendons. The micro: tissue strain ratios were less than 1 for all tendons, demonstrating that strain is attenuated and the amount of strain attenuation is dependent on the applied strain. The micro-scale sliding increased with the applied strain for both plantaris and tail tendons, suggesting that load is transferred via relative shearing at the micro-scale. The Micro-scale sliding did not correlate with the applied strain in Achilles tendon. Dashed lines represent 95% confidence intervals.

### Micro-scale Strains and Tendon Damage Mechanisms

All three tendons had a similar micro-scale strain and sliding response (Fig 5). The micro-scale strain fully recovered at the End of the Rest period (Fig 5a-c); however, the micro-scale sliding did not fully recover (Fig 5d-f). In addition, the non-recoverable sliding in percentage was strain-dependent for both plantaris and tail tendons (Fig 5g, i, p<0.005), and not for the Achilles tendon. Interestingly, the overall magnitudes of micro-scale sliding in degrees were similar for the plantaris and Achilles tendons loaded to 14% strain (Fig 5d-e), and no statistical difference of non-recoverable sliding in degrees was observed between the plantaris and Achilles tendons (Fig S-6, p>0.9). Note that this is the only place that we observed a similarly between the plantaris and Achilles tendons. Still, the non-recoverable sliding in percentage was much lower for the plantaris tendon because the micro-scale sliding at the End of SR was lower for the Achilles tendon (Fig 5h). The non-recoverable sliding in degrees and a close up of micro-scale sliding response of Achilles tendon are included as a supplemental material (Fig S-7).

**Fig 5:**
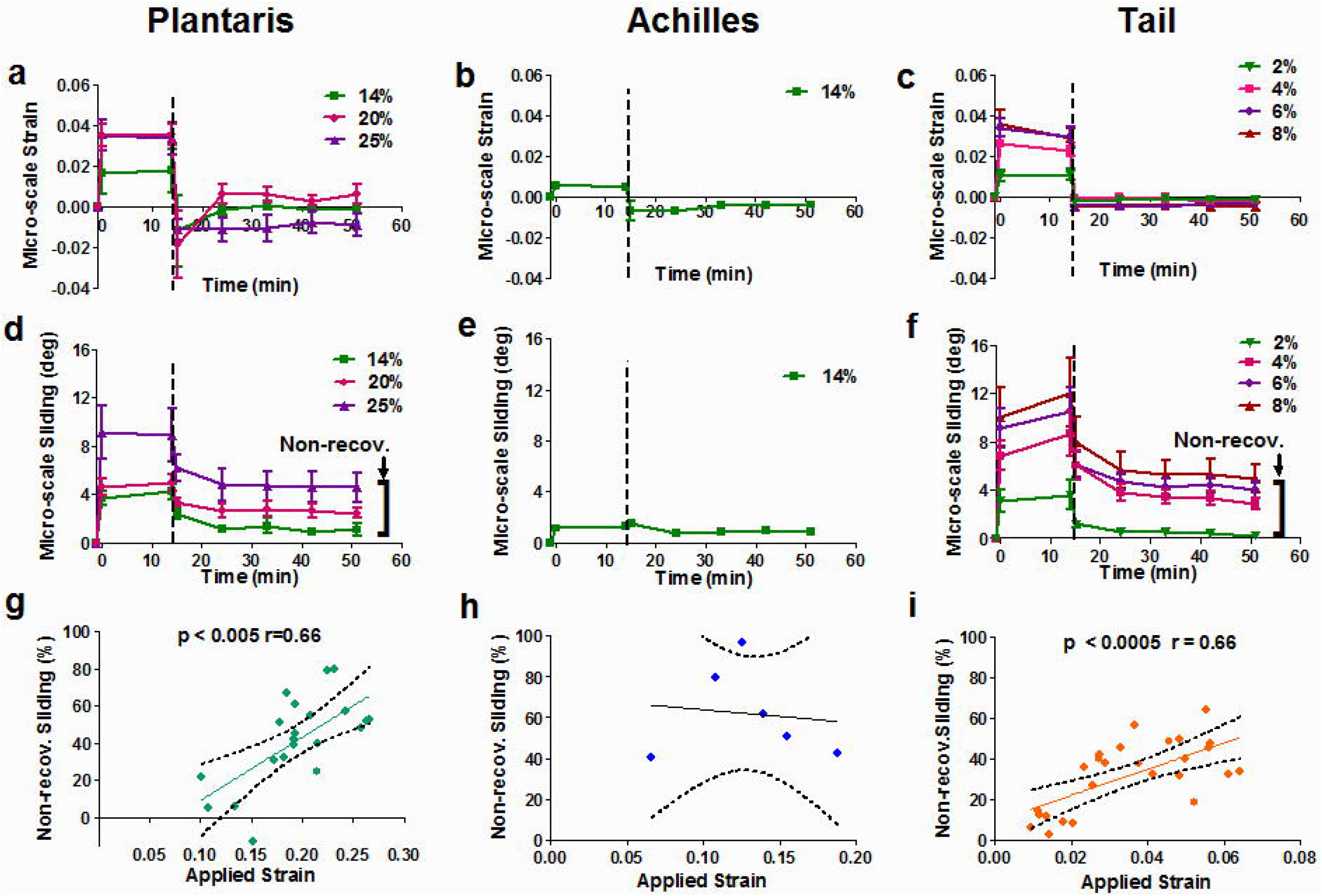
Overall micro-scale deformations. a)-c) For all three tendons, both the micro-scale strain and sliding had similar overall micro-scale deformation, where the micro-scale strain fully recovered at the End of the Rest Period. d)-f) On the other hand, the micro-scale sliding was not fully recoverable, suggesting the damage is observable by the micro-scale sliding. g)-i) The non-recoverable sliding was correlated with the applied strain for the plantaris and tail tendons but not for the Achilles tendon. Error bars are SEM and dashed lines represent 95% confidence intervals.

To compare across all three tendons, we evaluated the multi-scale mechanics with the normalized applied strain (Fig 6), calculated as the applied strain divided by the average peak strain during Ramp to Failure (Fig S-3). The directions of the correlations for the micro: tissue strain ratio (Fig 6a), micro-scale sliding during loading (Fig 6b), and non-recoverable sliding during recovery (Fig 6c) were the same for the plantaris and tail tendons. Indeed, the percentage of non-recoverable sliding overlapped between the plantaris and tail tendons (Fig 6c). The direction of correlation for the micro-scale deformation and non-recoverable sliding in percentage between plantaris and tail tendons suggest that their loading and damage mechanisms are similar.

**Fig 6:**
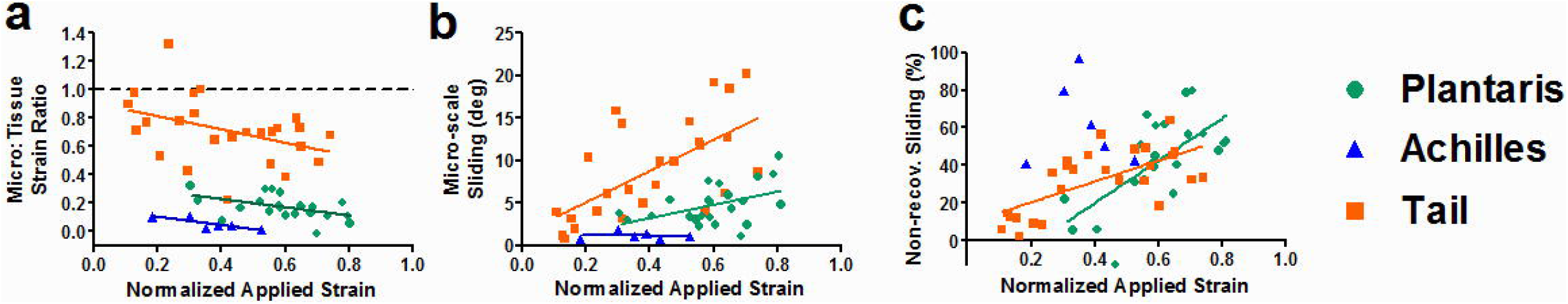
Normalized applied strain. a) The applied strain was normalized to plot on the same x-axis, and the directions of correlation for the micro: tissue strain ratio were the same for three tendons. b) The directions of micro-scale sliding between the plantaris and tail tendons were also the same but not for the Achilles tendon. c) The non-recoverable sliding directions and % magnitudes between plantaris and tail were notably comparable. Dashed lines represent 95% confidence intervals.

### Multi-scale Damage Mechanism Relationship between Tissue-scale and Micro-scale Parameters

Finally, we evaluated the relationship between the micro-scale damage (non-recoverable sliding) and tissue-scale mechanical properties. The transition strain was correlated with the non-recoverable sliding for both plantaris and tail tendons (Fig 7a-b). The transition strain is an indirect measurement of tissue elongation, and thus this suggests that an increase in tissue elongation is related to the irreversible microstructure damage. Although we observed correlations between non-recoverable sliding and tissue-scale inflection point strain and modulus in the tail tendon, these did not correlate in the plantaris tendon (Fig 7c-f).

**Fig 7:**
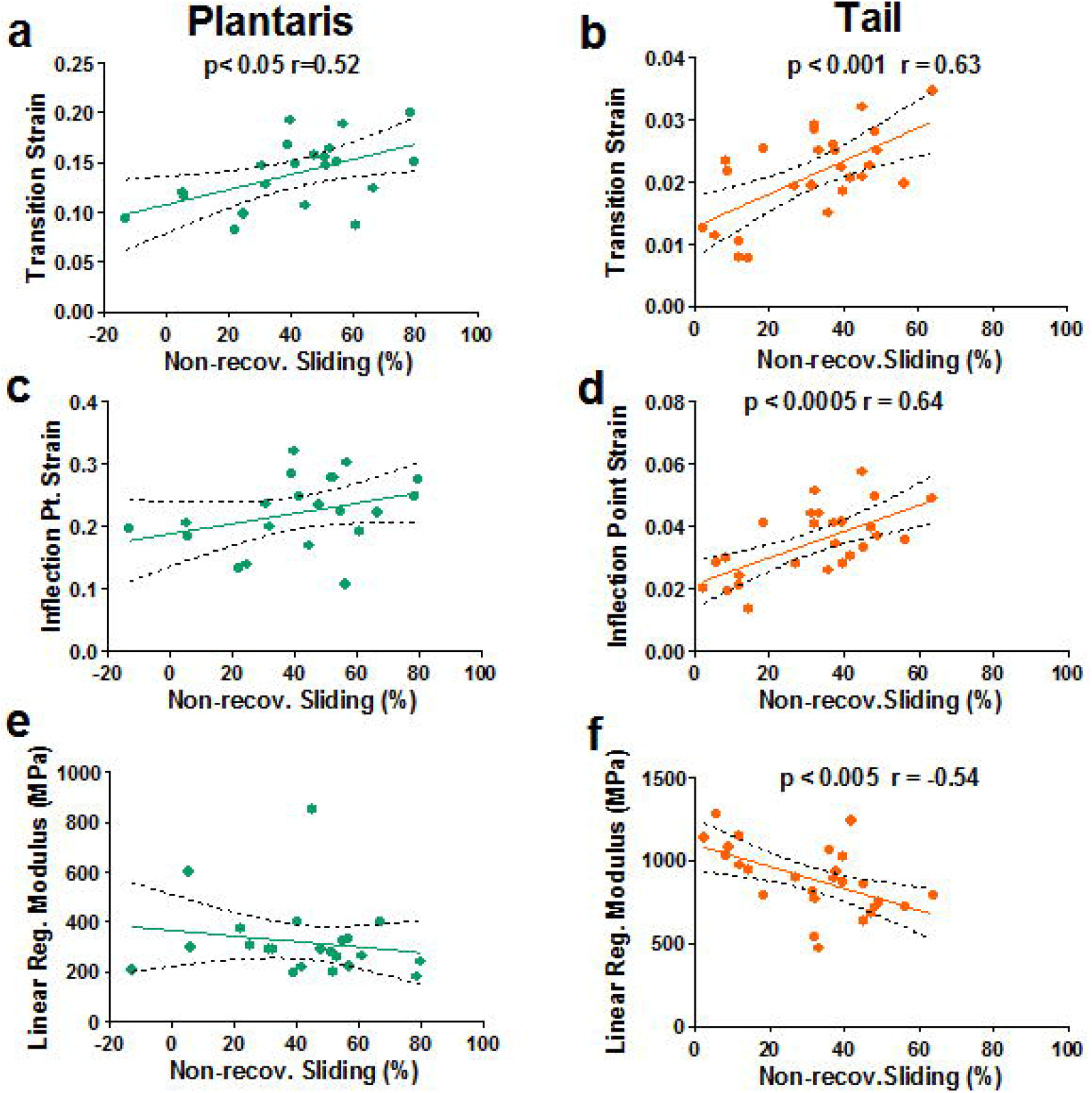
Relationship between tissue- and micro-scale parameters. a)-b) The transition strain was correlated with non-recoverable sliding. The transition strain is an indirect measurement of tissue elongation. and this suggests that an increase in tissue elongation is due to irreversible microstructure damage. c)-f) The inflection point strain and linear region modulus were only correlated for the tail tendon and not for the plantaris tendon. The correlation between micro-to tissue-scale parameter demonstrates that micro-scale sliding may be the damage mechanisms in both tendons. Dashed lines represent 95% confidence intervals.

## Discussion

This study investigated the normal load transfer and damage mechanisms of load-bearing tendons, the plantaris and Achilles tendons, across multiple length scales and showed that the mechanisms in the plantaris tendon are similar to those in tail tendon. Specifically, at the tissue-scale, damage was observed in the mechanical properties and was strain-dependent (Fig 3). At the micro-scale, the strain was highly attenuated in all tendons. The micro-scale sliding was correlated with applied strain, demonstrating that load was transferred via micro-scale sliding (shear) in the plantaris and tail tendons (Fig 4). The micro-scale sliding was non-recoverable and strain-dependent (Fig 5), and correlated with tissue-scale properties (Fig 7), suggesting the non-recoverable sliding is an initial mechanism of tendon damage (Fig 5). In summary, micro-scale sliding is likely responsible for both load transfer and damage mechanisms in load-bearing tendons.

### Damage Mechanisms of Functionally Distinct Tendons

While this study showed that micro-scale sliding is the mechanism of damage for both load-bearing rat tendons and non-load-bearing tail tendon, previous findings have suggested that functionally distinct tendons may not have the same damage mechanisms. This can be attributed to two main factors: 1) differences in structure and composition between larger and smaller animal tendons, and 2) differences in prescribed loading conditions. First, differences between low- and high-stress bearing tendons in equine and bovine tendons were observed in tendon mechanics,^37,38^ fascicle and interfascicular mechanics,^37,39^ interfascicular matrix composition,^40^ cross-link density,^38^ and fibril deformation.^38,41^ These observations suggest that damage mechanisms may depend on the mechanical function of a tendon, contrasting our findings; however, the difference in mechanisms may be explained through the presence/absence of fascicular and interfascicular structures. While larger animal tendons have fascicles and interfascicular matrix, rat tendons lack fascicles that are structurally and mechanically equivalent to those observed in larger animals.^10^ To support this notion, prior studies observed a significant difference in fascicle and sliding deformation of interfascicular matrix between functionally distinct tendons^37,39^, and thus fascicle and interfascicular matrix mechanics may dictate the damage mechanisms of functionally distinct tendons. The absence of fascicular and interfascicular structures in rat tendons likely explains the observed common damage mechanisms between functionally distinct tendons in rat.

Second, while this study showed non-recoverable sliding is the mechanism for damage, other recent studies have demonstrated that fibrils rupture under high magnitude and/or cyclic loading.^18,38,41^ Here, we observed a complete recovery of micro-scale strain, suggesting that fibrils did not rupture under the applied loading conditions. This challenges previous findings where the damage was observed in the form of collagen kinking, discontinuities, and denaturation.^13,16,18,38^ This disagreement may be attributed to the notion that damage observed as micro-scale sliding <u>may precede</u> the damage observed as micro-scale strain and fibril rupture.^4,42–44^ It is possible that micro-scale sliding damage precedes, regardless of the tendon type, and that damage related to fibril rupture takes place later, where this process may depend on the tendon type and loading protocol. It is likely that damage in the non-recoverable micro-scale strain will be observed in rat tendons after high magnitude and/or cyclic loading in forms of collagen kinking, discontinuities, and denaturation.

### Unique Features of the Plantaris and Achilles Tendons

The plantaris and Achilles tendons have unique mechanical features that can be considered in the context of their tissue-scale structure.^10^ The plantaris tendon has a single muscle-tendon junction and bone-tendon insertion, while the Achilles tendon has three muscle-tendon junctions and a single bone-tendon insertion.^10^ This complex structure of Achilles tendon likely contributes to the observed mechanics. The observed *σ-ε* response showed that the Achilles tendon bears high strain but low stress, and this is likely due to a combination of both micro-scale and inter-tendon slidings.^30^ The inter-tendon sliding, a sliding between sub-tendons from each muscle in Achilles tendon, has been speculated to contribute to low strain transfer from tissue-to micro-scale and inhomogeneous deformation.^40,41,47^ Notably, the tendon *σ-ε* response observed in rat plantaris (high stress) and Achilles (high strain and low stress) and inhomogeneous deformation have also been observed in human^30,31,36,45,46^ and mouse.^47^ In addition, the mechanical properties of rat Achilles tendons quantified in other studies were comparable to our mechanical properties.^48,49^ The complex structure, *σ-ε* response, lower strain transfer, and inhomogeneous deformation observed in this study and previous findings suggest that Achilles tendon should be recognized as a specialized tendon, and the structural complexity of Achilles tendon should be carefully considered. Based on the structure and *σ-ε* response, we recommend selecting the rat plantaris tendon over the Achilles tendon as a small animal model system when studying load-bearing tendons.

### Strain Attenuation and Micro-Scale Strain of Native Tissue

This study demonstrated that strain is highly attenuated in rat tendons, and since the micro-scale strain is a good indicative measure for cell strain^50^, native cells in tendon will likely experience a small relative amount of strain, regardless of the applied strain. Mechanical stimulation is a crucial aspect of maintaining homeostasis of tendon cells, and thus strain transfer from the tissue-to micro-scale structure is important.^50^ Interestingly, while the plantaris and Achilles tendons were loaded to an applied strain that was about three times higher than the tail tendon, the magnitudes (in degree) of micro-scale strain were comparable, likely due to the load transfer mechanism of micro-scale sliding.^14^ These observations are consistent with the magnitude of micro-scale strain measured in other aligned fibrous tissues such as meniscus and annulus fibrosus,^50^ suggesting that cells in the native environment may only require low strain to maintain homeostasis in a healthy state. This finding has important implications for mechanotransduction and the design of tissue engineered constructs.

## Conclusion

In summary, this study determined the multi-scale load transfer and damage mechanisms in the rat plantaris, a load-bearing tendon, by demonstrating that the micro-scale sliding is a key component in both mechanisms. These mechanisms are similar to those previously quantified in the tail tendon. Namely, the micro-scale sliding correlated with the applied strain measured at the End of the Initial Ramp, suggesting that micro-scale sliding contributes to load transfer in tendon. In addition, while the micro-scale strain fully recovered, the micro-scale sliding was non-recoverable and strain-dependent, and correlated with tissue-scale mechanical parameter. When the applied strain was normalized, the percentage magnitude of non-recoverable sliding was similar between the plantaris and tail tendons. Achilles tendon demonstrated some of the mechanical responses observed in plantaris and tail tendons, yet the results were inconclusive due to its complex structure. Thus, micro-scale sliding is responsible for both loading and damage mechanisms in load bearing plantaris tendon.

## Acknowledgment

This research was supported and made possible by the National Institutes of Health grant No. R01 EB002425, the National Institute of General Medical Sciences grant No. P20 GM103446, National Institute of General Medical Sciences grant No. S10 RR027273, the National Science Foundation grant No. IIA-1301765, and the State of Delaware. We thank the Bioimaging Center at the Delaware Biotechnology Institute and Dr. Jeff Caplan for supporting data acquisition. We thank Francis Karani for providing rats for tendon.

Authors Contribution
A.H.L. and D.M.E. designed experiments. A.H.L. performed experiments and analyzed data. A.H.L. and D.M.E. wrote the manuscript.

